# Cross-sectional genomic perspective of epidemic waves of SARS-CoV-2: a pan India study

**DOI:** 10.1101/2021.08.11.455899

**Authors:** Sanjeet Kumar, Kanika Bansal

## Abstract

COVID-19 has posed unforeseen circumstances and throttled major economies worldwide. India has witnessed two waves affecting around 31 million people representing 16% of the cases globally. To date, the epidemic waves have not been comprehensively investigated to understand pandemic progress in India. In the present study, we aim for a cross-sectional analysis since its first incidence up to 26th July 2021. We have performed the pan Indian evolutionary study using 20,086 high-quality complete genomes of SARS-CoV-2. Based on the number of cases reported and mutation rates, we could divide the Indian epidemic into seven different phases. First, three phases constituting the pre-first wave had a very less average mutation rate (<11), which increased in the first wave to 17 and then doubled in the second wave (~34). In accordance with the mutation rate, variants of concern (alpha, beta, gamma and delta) and interest (eta and kappa) also started appearing in the first wave (1.5% of the genomes), which dominated the second (~96% of genomes) and post-second wave (100% of genomes) phases. Whole genome-based phylogeny could demarcate the post-first wave isolates from previous ones by the point of diversification leading to incidences of VOCs and VOIs in India. Nation-wide mutational analysis depicted more than 0.5 million events with four major mutations in ~97% of the total 20,086 genomes in the study. These included two mutations in coding (spike (D614G) and NSP 12b (P314L) of RNA dependent RNA polymerase), one silent mutation (NSP3 F106F) and one extragenic mutation (5’ UTR 241). Large scale genome-wide mutational analysis is crucial in expanding knowledge on evolution of deadly variants of SARS-CoV-2 and timely management of the pandemic.

## Introduction

Coronavirus represents a large family of RNA viruses causing upper and lower respiratory tract infections to humans ranging from mild to lethal. Previously reported outbreaks of coronaviruses causing great public health threats include Severe Acute Respiratory Syndrome (SARS) and Middle East Respiratory Syndrome (MERS) (Peiris, Guan et al. 2004, Memish, Cotten et al. 2014). In late December 2019, the ongoing outbreak caused by novel coronavirus epi-centered in Hubei province of People’s Republic of China (Chen, Liu et al. 2020, Wu, Zhao et al. 2020). Patients were epidemiologically linked to a wet animal and seafood wholesale market in Wuhan (Bogoch, Watts et al. 2020, Lu, Stratton et al. 2020). On the basis of phylogeny and taxonomic analysis Coronavirus Study Group of International Committee on Taxonomy of Viruses recognized this as a sister to SARS-CoV (Peiris, Guan et al. 2004) and named it as SARS-CoV-2 (Gorbalenya, Baker et al. 2020). SARS-CoV-2 has the largest genome (26-4 to 31.7 kb) among all known RNA viruses with variable GC content ranging from 32% to 43% (Woo, Huang et al. 2010). On 30^th^ January 2020, a global health emergency was declared by the WHO Emergency Committee. Due to its substantial human to human transmissions, it has spread to many countries and till now it has affected more than 195 million people with more than 4 million casualties worldwide (Lu, Zhao et al. 2020, Worldometer 2020).

First case of SARS-CoV-2 from India was reported in Kerala during 27-31 January, 2020 in individuals with travel history of Wuhan, China (Andrews, Areekal et al. 2020). In order to contain further spread of SARS-CoV-2 strict actions were imposed including international travel restrictions from the affected countries. However, a surge of COVID-19 cases began in the month of March 2020 along with several local transmission events (https://www.worldometers.info/coronavirus/country/india/; https://www.covid19india.org/). From 26th March to 11th May 2020 a nationwide lockdown was imposed to contain the nationwide spread. This was one of the most rigid lockdown restrictions in the world which resulted in a well-controlled infectivity rate in India (Maitra, Raghav et al. 2020, Mitra, Misra et al. 2020). After the unlocking despite the measures, India witnessed a surge in cases resulting in the first wave from July till December 2020. Nationwide first wave peaked at 93,732 on 17th September 2020 and six weeks later the toll had fallen by half. Since the middle of March 2021, a second wave has started in India which peaked at 3,91,236 cases per day by 8th May 2021. This wave witnessed a breakneck rise of COVID-19 cases which overwhelmed the healthcare system in the country. In the beginning, there appeared to be a lack of coordination between the state and central government of India, but soon the strict lockdown restrictions, proper identification of containment zones played a key role in the fall of the second wave. Second wave slowed down in record time of around three weeks, contrast to the first wave which lasted for several months. Second wave was then reported to end by June 2021. During both the waves, Delhi and Maharashtra were most badly affected states with several localized outbursts due to rampant community transmission. As of 26^th^ July 2021, India is reporting 30820 cases per day with total cases of 3,14,40,492 and the death toll has also reached 421,414. Yet, a nationwide third wave is also predicted which is concerning for the policy makers and public governance.

We have witnessed generation of unprecedented genomic resources of SARS-CoV-2 worldwide such as by UK Consortium (Gorbalenya 2020), African union (Salyer, Maeda et al. 2021), Indian SARS-CoV-2 Genomic Consortia (INSACOG) (Maitra, Raghav et al. 2020, Alai, Gujar et al. 2021) etc. GISAID (https://www.gisaid.org/), which is a global initiative for a public repository for the genomic data of SARS-COV-2 storage and analysis. Such a vast genomic resource has been investigated in detailed based on mutation in SARS-CoV-2 into various lineages (Rambaut, Holmes et al. 2020). On the basis of these studies, World Health Organisation have announced variants of concern (VOC) (alpha, beta, gamma and delta) and variants of interest (VOI) (eta, lota, kappa and lambda) (https://www.who.int/en/activities/tracking-SARS-CoV-2-variants/). These VOCs and VOIs are known to pose increased risk to the public health globally and will aid in monitoring the evolution of deadly variants worldwide.

In order to understand the genetic diversity, transmission and cure in human, several large-scale genome-based studies have been conducted (Bajaj and Purohit 2020, Helmy, Fawzy et al. 2020, Phan 2020, Salyer, Maeda et al. 2021, Yadav, Nyayanit et al. 2021). Pan India study of 1000 sequences across 10 states suggest the widespread presence of the several lineages of SARS-CoV-2 (Maitra, Raghav et al. 2020). Pan India sero survey suggested average seropositivity to be 10.14% (among 10,427 subjects) (Naushin, Sardana et al. 2021). Unfortunately, the spread of lineages across India and seropositivity rate is very complex due to the vast population and landmass. Genomic diversity of Indian isolates in comparison to the global lineages is supposed to evolve further which needs to be closely monitored (Alai, Gujar et al. 2021).

Till date pan India genome-based studies are focused on evolution of SARS-CoV-2 only upto first wave (Maitra Raghav et al. 2020, Alai, Gujar et al. 2021, Yadav, Nyayanit et al. 2021). However, since then India has witnessed devastating second wave with more than 0.4 million cases per day which is four times the cases reported during first wave (https://www.worldometers.info/coronavirus/country/india/; https://www.covid19india.org/). This created a lacuna in understanding the evolution of deadly variants of SARS-CoV-2. Since the first epidemic wave there is an upsurge in public genomic resources of SARS-CoV-2 from India which is available from global GISAID initiative. Current scenario provides scope of a crosssectional genome-based monitoring of the deadly variants across India. In the present study, we have analysed 20,086 high quality genomes to understand the dominance of VOCs and VOIs in the second wave. Amongst 0.5 million mutations we could identify four major mutation events in more than 19,300 genomes in spike, RNA dependent RNA polymerase and extragenic 5’UTR. Our analysis depicted single nucleotide transitions as the major player in the evolution of SARS-CoV-2 in India. Pan India study based on the mutation can open a gate way to understand the hotspots of mutation in SARS-CoV-2.

## Results and Discussion

### Evolutionary timeline of epidemic waves of SARS-CoV-2 in India

Since the first incidence of COVID-19 in India by the end of January 2020, the number of cases has increased abruptly twice, which we call first and second wave. In order to understand the peak and plateau of incidences, we have differentiated the time period from first incidence (January 2020) to 26th July 2021 in seven different phases (Figure 1 C). Here, phase I indicates the early days of infection i.e., introductory phase from January 2020 to 25th March 2020. Phase II refers to the nationwide lockdown from 26th March 2020 to 11th May 2020 which was imposed to contain the spread of COVID-19. Once the lockdown was relaxed the incidences of cases rose gradually, termed as pre-first wave period or phase III from 12th May 2020 to 31st June 2020. Phase IV or first wave of COVID-19 was demarcated for quite a long duration from 1st July 2020 to 31st December 2020. India had witnessed a peak incidence of 93,732 cases of COVID-19 on 17th September 2020 (https://www.worldometers.info/coronavirus/country/india/; https://www.covid19india.org/) during phase IV (Figure 1 A). With the gradual decrease in number of cases across India, phase V was demarcated from 1st January 2021 to 15th March 2021 as post-first wave or pre-second wave. Once again major leap in incidences was observed resulting in second wave which we refer as phase VI from 16th March 2021 to 30th June 2021. Strikingly, the rise and fall of incidences during the second wave (phase VI) was very steep as compared to the first wave (phase IV). Currently India is undergoing phase VII with drastic reduction in overall incidences from 1st July 2021 up to now.

**Figure 1:**
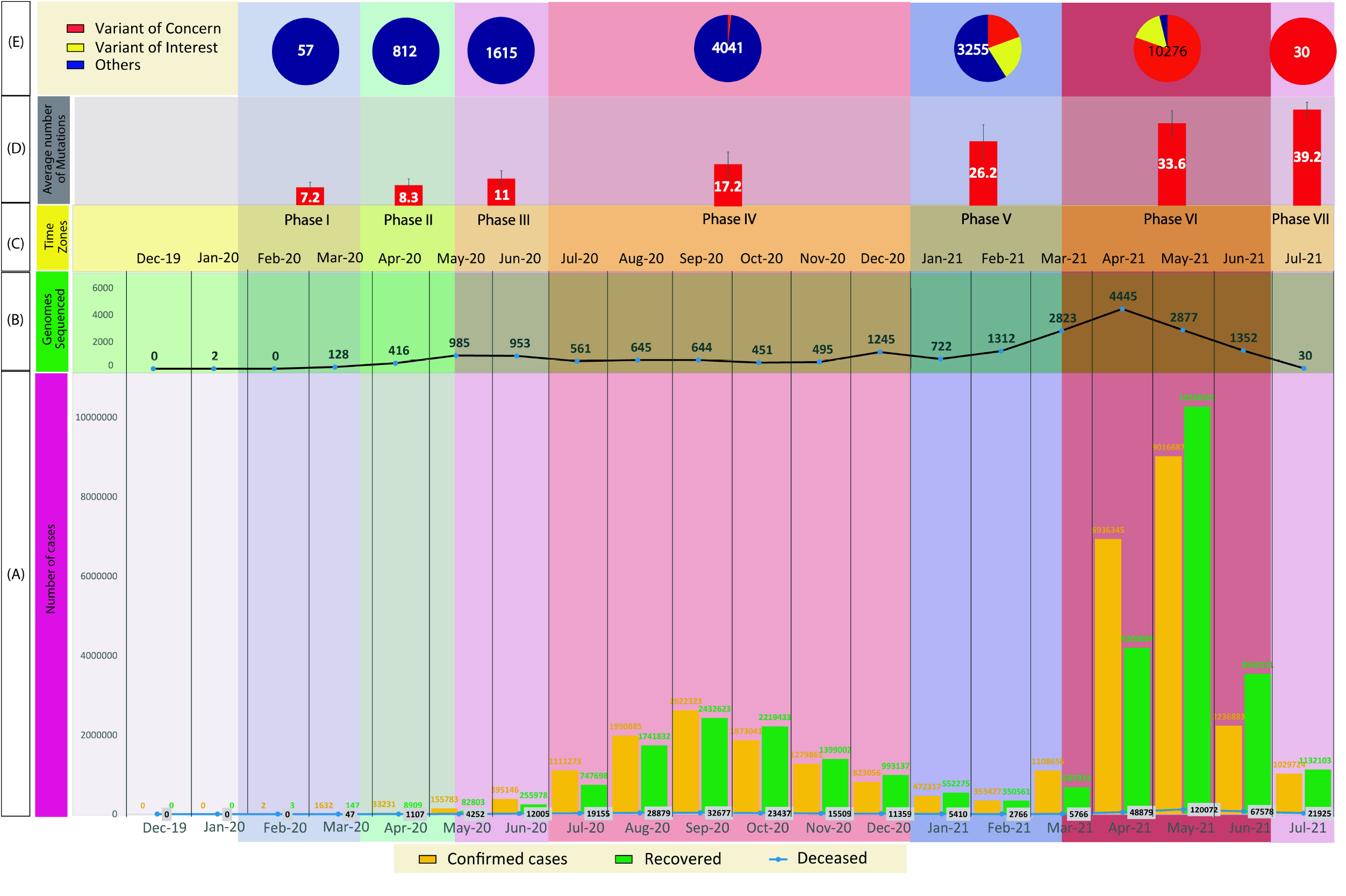
Pan Indian overview of SARS-CoV-2 across seven phases. **A).** Bar graph plot of number of confirmed cases (yellow), recovered (green) and deceased (black) during each month since their first incidence in India. **B).** Number of genome sequences submitted in the public repository of GISAID month wise since its first incidence. **C).** Seven phases of SARS-CoV-2 pandemic and their time zones are represented as phase I: introductory phase, phase II: nationwide lockdown, phase III: pre-first wave, phase IV: first wave, phase V: post-first wave/ pre-second wave, phase VI: second wave and phase VII: post-second wave. **D).** Average number of mutations during each phase. Standard deviation in the mutation rate is indicated by a vertical line. E). Distribution of variants of concern (VOC) and variants of interest (VOI) and other lineages defined in accordance with pangolin lineage. The total number of genomes included in each phase is marked in the center of the pie chart.

### Nation-wide phylogenetic network of SARS-CoV-2

India has witnessed two epidemic waves of SARS-CoV-2, the number of cases and genome resources have also increased accordingly. By the end of the first wave, India reported 6,525 high quality genomes which increased by another 13,531 by the end of the second wave and still continuing (Figure 1 B). Overall, upto 26th July 2021, India has reported 3,17,25,450 cases and 20,086 high quality genomes (Supplementary Figure 1 and Supplementary Table 1) (https://www.gisaid.org/; https://www.worldometers.info/coronavirus/country/india/; https://www.covid19india.org/).

Pan India phylogeny based on whole genome sequences (n=20,086) have revealed major lineages of SARS-CoV-2 in India (Table 1). It clearly demarcated post-first wave isolates (phase V, VI and VII) from the earlier isolates (phase I, II, III and IV) (Figure 2). Post-first wave represents recent introduction of deadly variants in the Indian population of SARS-CoV-2 (Figure 1 E and Figure 2). We could identify the clade representing point of diversification (marked as red dot in figure 2) as the major event in evolution of SARS-CoV-2.

**Table 1:** Lineage distribution amongst the genomes of different time zones

|  | Phase I (Introductory Phase) | Phase II (Nationwide lockdown) | Phase III (Pre-first wave) | Phase IV (First wave) | Phase V (Post-first wave/ Pre-second wave) | Phase VI (Second wave) | Phase VII (Post-second wave) |
| --- | --- | --- | --- | --- | --- | --- | --- |
| PANGOLIN LINEAGE |  |  |  |  |  |  |  |
| A | 4 | 17 | 15 | 6 | - | - | - |
| A.1 | 2 | - | - | - | - | - | - |
| A.2 | - | 1 | - | 1 | 4 | 3 | - |
| A.7 | - | 12 | - | - | - | - | - |
| A.9 | - | 16 | 6 | 1 | - | - | - |
| AE.1 | - | - | - | - | 1 | 1 | - |
| AE.2 | - | - | - | 1 | - | - | - |
| AM.3 | - | - | - | 2 | 1 | - | - |
| AY.1 | - | - | - | - | - | 6 | - |
| AY.3 | - | - | - | - | - | 2 | 1 |
| B | 1 | 5 | 7 | 12 | 1 | - | - |
| B.1 | 31 | 476 | 1520 | 4002 | 3239 | 10258 | 29 |
| B.4 | 13 | 3 | 4 | 5 | - | - | - |
| B.53 | 1 | - | - | - | - | - | - |
| B.6 | 5 | 282 | 63 | 9 | 3 | 1 | - |
| C.36 | - | - | - | 1 | 3 | 3 | - |
| L.3 | - | - | - | - | 1 | - | - |
| P.1 | - | - | - | - | - | 1 | - |
| P.2 | - | - | - | 1 | - | - | - |
| R.1 | - | - | - | - | 2 | - | - |
| GISAID LINEAGE |  |  |  |  |  |  |  |
| G | 14 | 214 | 372 | 522 | 1316 | 8382 | 13 |
| GH | 8 | 103 | 439 | 1576 | 849 | 178 | - |
| GK | - | - | - | - | - | 777 | 17 |
| GR | 9 | 154 | 675 | 1847 | 739 | 426 | - |
| GRY | - | - | - | 23 | 331 | 460 | - |
| GV | - | - | - | 5 | 3 | 6 | - |
| L | 3 | 10 | 3 | 6 | 1 | - | - |
| O | 15 | 288 | 103 | 54 | 12 | 44 | - |
| S | 6 | 42 | 23 | 8 | 4 | 3 | - |
| V | 2 | 1 | - | - | - | - | - |

**Figure 2:**
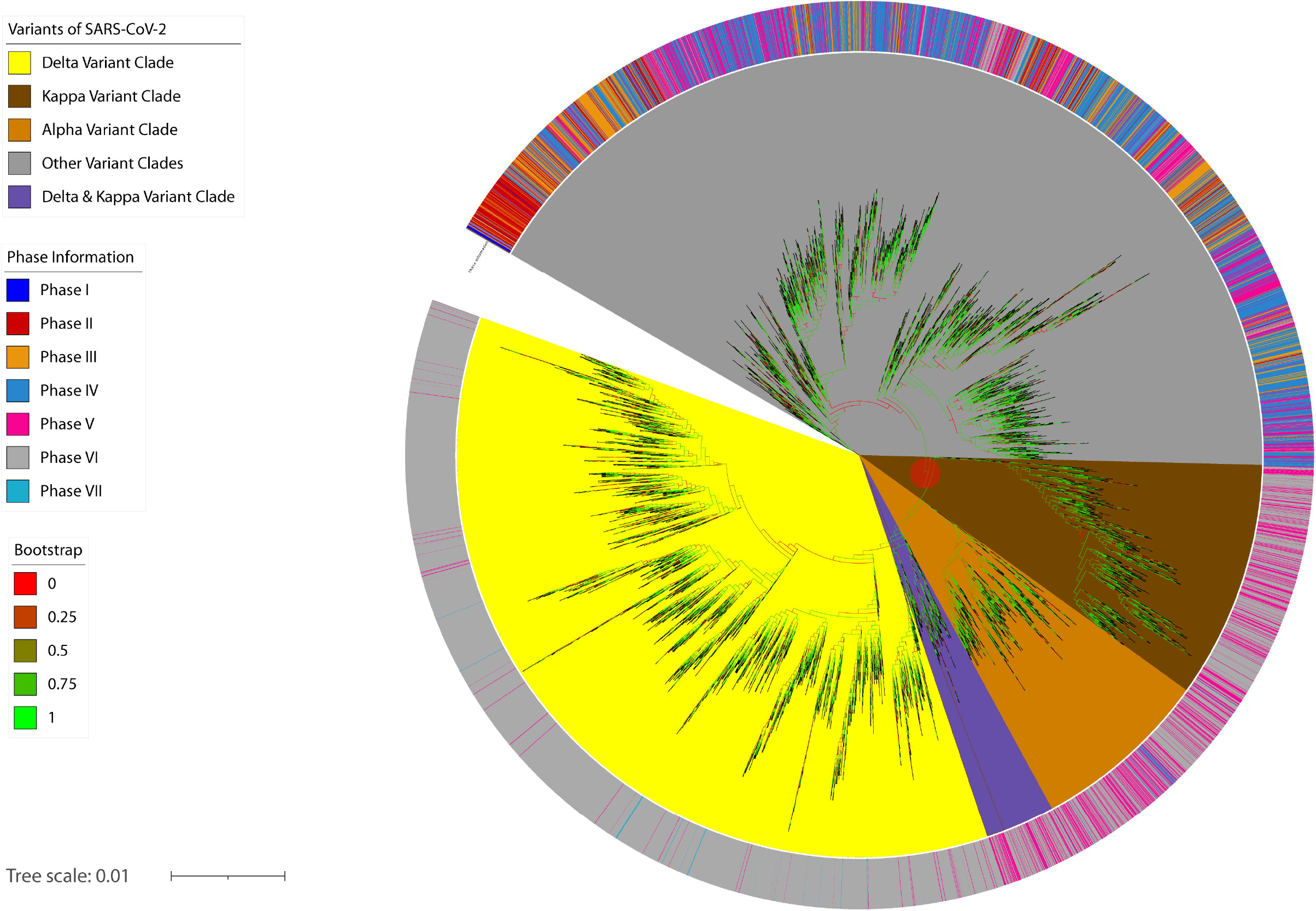
Pan India whole genome-based phylogeny of SARS-CoV-2. Here, bootstrap values are represented in the color range from 0 (minimum value clade marked with red) to 1 (maximum value clade marked with green). Clades representing variants of SARS-CoV-2 (delta, kappa, alpha etc.) are marked with respective colors as indicated. Isolates reported in different phases are marked with the color strip against their respective leaf in the phylogenetic tree. The point of diversification is indicated by a red dot in the phylogenetic tree.

In India second wave outbreak has been a record-buster with impulsive rise in infection from 0.7% to 1.06% of the total population within two months. During this time maximum per day cases reported were around 0.4 million which is way beyond than recorded worldwide. Such a widespread of virus imparts opportunities of mutation and provides fitness to its variants. Post-first wave 83% of the genomes could be linked to the deadly variants (VOCs and VOIs) (Table 3 and Supplementary Table 2). Among them, delta, alpha and kappa were dominant in phases V, VI and VII (Figure 2).

### Mutations driving emergence of VOCs and VOIs

In addition to the confirmed cases, seven phases of the pandemic in India can also be distinguished on the basis of mutations detected in the rapidly evolving virus. For instance, the average mutation detected before the first wave was less than 11 which elevated to 17.2 and 33.6 in the first and second waves respectively (Figure 1 D). Hence, average mutations were doubled in the second wave as compared to the first wave and is still in the rising trend.

Strikingly, in accordance to the phylogeny, rise in mutations is directly correlated with the emergence of deadly variants (VOCs and VOIs) in India. For instance, based on genomic analysis, these deadly variants were reported during the first wave, yet cases due to them started alarmingly incriminating only after the first wave (Figure 1 E). However, the second wave was dominated by these deadly viruses and the post-second wave is hauntingly related to these deadly variants only (this is based on just 30 genomes available till 26th July for phase VII).

According to the pangolin lineages out of 20,086 stains used in the present study, 7421 were delta variants (B.1.617.2, AY.1, AY.2 and AY.3), 95 were beta (B.1.351) and 1 gamma (P.1) constituting VOCs and 42 eta (B.1.525), 2300 kappa (B.1.617.1) constituting VOIs (Table 2). None of the VOCs or VOIs were reported in India upto phase III i.e., before first wave. These started appearing only in first wave (phase IV). Alpha, delta and kappa variants were detected in 1.5% of the phase IV genomes analysed. Phase V had 41 % of the genomes of alpha, beta, delta, eta and kappa variants. Further, second wave or phase VI had 96 % of the cases by these deadly variants including alpha, beta, gamma, delta, eta and kappa. While post-second wave is found to be dominated by delta variants only (based on only 30 genomes sequenced till 26th July 2021) (Table 1). Lota and lambda were not detected in India based on the genome sequencing data available till date.

**Table 2:** Distribution of variant of concern (VOC) and variant of interest (VOI) down the timeline in India

|  | High quality genomes | VOC (alpha + beta + gamma + delta) | VOI (eta + iota + kappa + lambda) |
| --- | --- | --- | --- |
| Phase I (Introductory phase) | 57 | 0 | 0 |
| Phase II (Nation-wide lockdown phase) | 812 | 0 | 0 |
| Phase III (Pre-first wave) | 1615 | 0 | 0 |
| Phase IV (First wave) | 4041 | 56 (55 + 0 + 0 + 1) | 7 (0 + 0 + 7 + 0) |
| Phase V (Pre-second or post-first wave) | 3255 | 632 (509 + 26 + 0 + 97) | 709 (6 + 0 + 703 + 0) |
| Phase VI (Second wave) | 10276 | 8259 (896 + 69 + 1 + 7293) | 1626 (36 + 0 + 1590 + 0) |
| Phase VII (Post-second wave) | 30 | 30 (0 + 0 + 0 + 30) | 0 (0 + 0 + 0 + 0) |

Further, nation-wide mutational analysis depicted more than 0.5 million mutations amongst the Indian SARS-CoV-2 population (Supplementary Table 3) from seven different phases (Supplementary Figure 2). Out of which we have looked at the top twenty prevalent mutations (Table 3). Here, top twenty mutations were basically found in spike, RNA dependent RNA polymerase, nucleocapsid, ORF3, ORF7 and extragenic regions (5’ UTR and 3’ UTR). Interestingly, there were four widespread mutations in ~97% of the genomes analysed essentially representing all over India since its emergence in the country. Two mutations in protein coding region i.e., D614G in spike and P314L in NSP 12b; one extragenic mutation at 241 position of 5’UTR and one silent mutation at F106 of NSP3 (Figure 3, Table 3). Such prevalent non-synonymous or silent mutations in spike protein and rdrp and 5’ UTR may likely to improve pathogenicity of the virus, in evasion from host immune system and risk of human-to-human transmissions etc.

**Table 3:** Top twenty mutations amongst the pan Indian SARS-CoV-2 isolates.

| Nucleotide mutation | Protein mutation | Number of strains | Gene | Mutation type | Annotation |
| --- | --- | --- | --- | --- | --- |
| A23403G | D614G | 19571 | Spike | Non-synonymous SNP | Spike |
| C3037T | F106F | 19527 | NSP3 | Synonymous mutation | Predicted phosphoesterase, papain-like proteinase |
| C241T | - | 19583 | 5'UTR | Extragenic | NA |
| C14408T | P314L | 19353 | NSP 12b | Non-synonymous SNP | RNA-dependent RNA polymerase, post-ribosomal frameshift |
| G210T | - | 9985 | 5'UTR | Extragenic | NA |
| C23604G | P681R | 9887 | Spike | Non-synonymous SNP | Spike |
| G29402T | D377Y | 9878 | N | Non-synonymous SNP | Nucleocapsid protein |
| T22917G | L452R | 9875 | Spike | Non-synonymous SNP | Spike |
| G29742T | - | 9836 | 3'UTR | Extragenic | NA |
| G28881T | R203M | 9835 | N | Non-synonymous SNP | Nucleocapsid protein |
| T27638C | V82A | 9813 | ORF7a | Non-synonymous SNP | ORF7a protein |
| C25469T | S26L | 9713 | ORF3a | Non-synonymous SNP | ORF3a protein |
| C21618G | T19R | 7473 | Spike | Non-synonymous SNP | Spike |
| T26767C | I82T | 7429 | M | Non-synonymous SNP | Membrane |
| C27752T | T120I | 7397 | ORF7a | Non-synonymous SNP | ORF7a protein |
| C22995A | T478K | 7376 | Spike | Non-synonymous SNP | Spike |
| A28461G | D63G | 6999 | N | Non-synonymous SNP | Nucleocapsid protein |
| G15451A | G662S | 6785 | NSP12b | Non-synonymous SNP | RNA-dependent RNA polymerase, post-ribosomal frameshift |
| C16466T | P77L | 6762 | NSP13 | Non-synonymous SNP | Helicase |
| G24410A | D950N | 5470 | Spike | Non-synonymous SNP | Spike |

**Figure 3:**
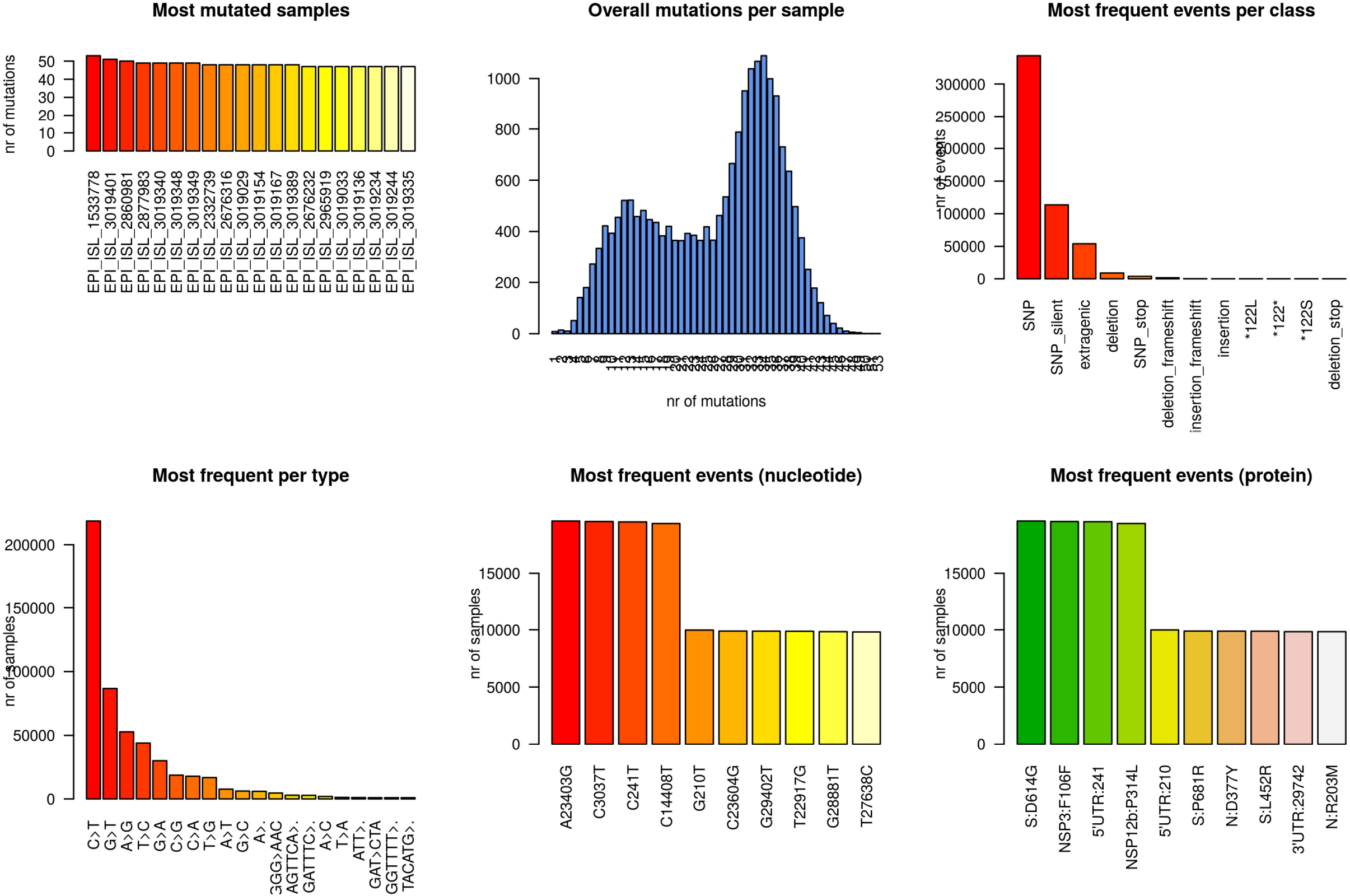
Pan India mutation analysis. Six panel image displays the most mutated samples, overall mutations per samples, most frequent events per class of mutation category, changes of nucleotide per type, nucleotide wise most frequent events and protein level most frequent events for 20,086 genomes used in the study.

## Methods

### Procurement of SARS-CoV-2 genome from public repository

We have considered 20,086 high quality genomes from India encompassing the country’s length and breadth. The source of the genomes we considered in this study is the EpiCoV database maintained under GISAID initiative (https://www.gisaid.org/). A detailed state-wise information is provided in supplementary information (Supplementary Table 1) indicating patients details such as geographical location, age, sex, pangolin lineage, GISAID lineage etc. First strain reported from Wuhan (China) NC_045512.2 (https://www.ncbi.nlm.nih.gov/genome/?term=NC_045512.2) was taken as a reference strain for all the analysis in the study. We have considered all high-quality complete genomes submitted until 26th of July 2021.

### Phylogenetic analysis

In order to obtain a pan India phylogeny of SARS-CoV-2, we have used all high quality genomes (n=20,086) available by 26th July 2021 in the public repository of GISAID (https://www.gisaid.org/). Multiple sequencing alignment (MSA) was performed using (MAFFT v7.467) (Nakamura, Yamada et al. 2018) (https://mafft.cbrc.jp/alignment/software/) which is based on fast fourier transform keeping NC 045512.2 (https://www.ncbi.nlm.nih.gov/genome/?term=NC_045512.2) (Wuhan-Hu-1) strain as reference. MSA obtained was used for the phylogenomic tree generation using fasttree v2.1.8 with double precision (Price, Dehal et al. 2010) with gamma time reversal method (gtr). Visualization of the phylogenomic tree was performed using a web server of iTOL v6 (https://itol.embl.de/) (Letunic and Bork 2019). The isolates were marked in accordance with their phase of isolation and pangolin lineage (Rambaut, Holmes et al. 2020).

### Pan India mutation analysis across different phases

In order to understand the rate of mutation across the genome procured for different phases (seven), we have first aligned all the sequences (n=20086) against NC 045512.2 (https://www.ncbi.nlm.nih.gov/genome/?term=NC_045512.2) (Wuhan-Hu-1) strain (reference strain) using nucmer v3.1 (Delcher, Phillippy et al. 2002). Also, we have separately aligned all the strains from several phases against the reference sequence. In order to translate all the alignments scores into mutational events, we have implemented a well-documented method earlier described by Mecatelli and Giorgi (Mercatelli and Giorgi 2020). This approach uses a gff3 annotation file and reference sequence of NC_045512.2 to extract the genomic coordinates of SARS-CoV-2 proteins. R library package seqinr (https://cran.r-project.org/web/packages/seqinr/index.html) and biostring package of bioconductor (https://bioconductor.org/packages/release/bioc/html/Biostrings.html) was used to get the list of mutational events in terms of nucleotide and protein. Frequency and rate of mutation per sample was also obtained for all samples and across each phase. Overall number of mutations, coordinates of mutations with respect to the reference strain were also calculated using same R script.

## Acknowledgment

Authors whole heartedly acknowledge motivation and advice from Dr. Prabhu B. Patil – CSIR-Institute of Microbial Technology. We do gratefully acknowledge GISAID for sharing the genomic sequences in public domain and several contributors of SARS-CoV-2 genomic data. We would also like to convey our acknowledgement to Government of India for enriching the genomic resources through initiatives like INSACOG. Further, authors would like to acknowledge Ms. Anu Singh for tea time chit-chat over SARS-CoV-2.

## Author contribution

KB and SK have performed the data procurement and analysis. KB has conceived and drafted the manuscript by taking inputs from SK.

## Data availability

All the metadata files generated in this study can be accessed through https://figshare.com/s/0a81433867e6e6df2cec.

## Figure and Table legends

**Supplementary Figure 1: A).** Total number of cases reported state wise in India until 26th July 2021. **B).** Genome sequence data available from each state from India until 26th July 2021.

**Supplementary Figure 2:** Overview of the mutation reports during seven different phases.

**Supplementary Table 1:** Metadata of all the genomes used in the present study.

**Supplementary Table 2:** Total VOCs and VOIs in Indian SARS-CoV-2 population.

**Supplementary Table 3:** Total mutations in Indian SARS-CoV-2 population.

