## Supplementary figures and images for "Cross-sectional genomic perspective of epidemic waves of SARS-CoV-2: a pan India study"

### Supplementary Figure 1

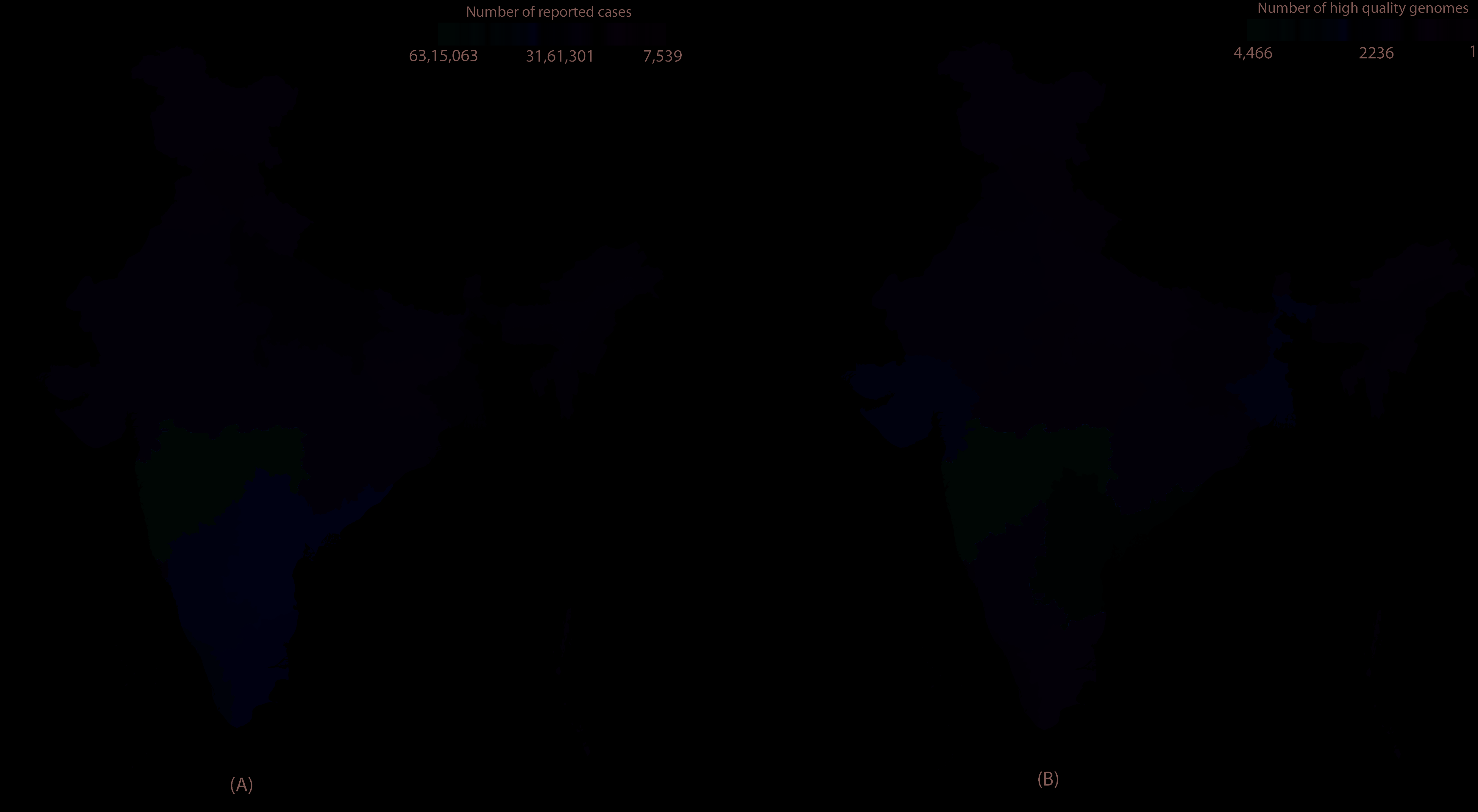

### Supplementary Figure 2

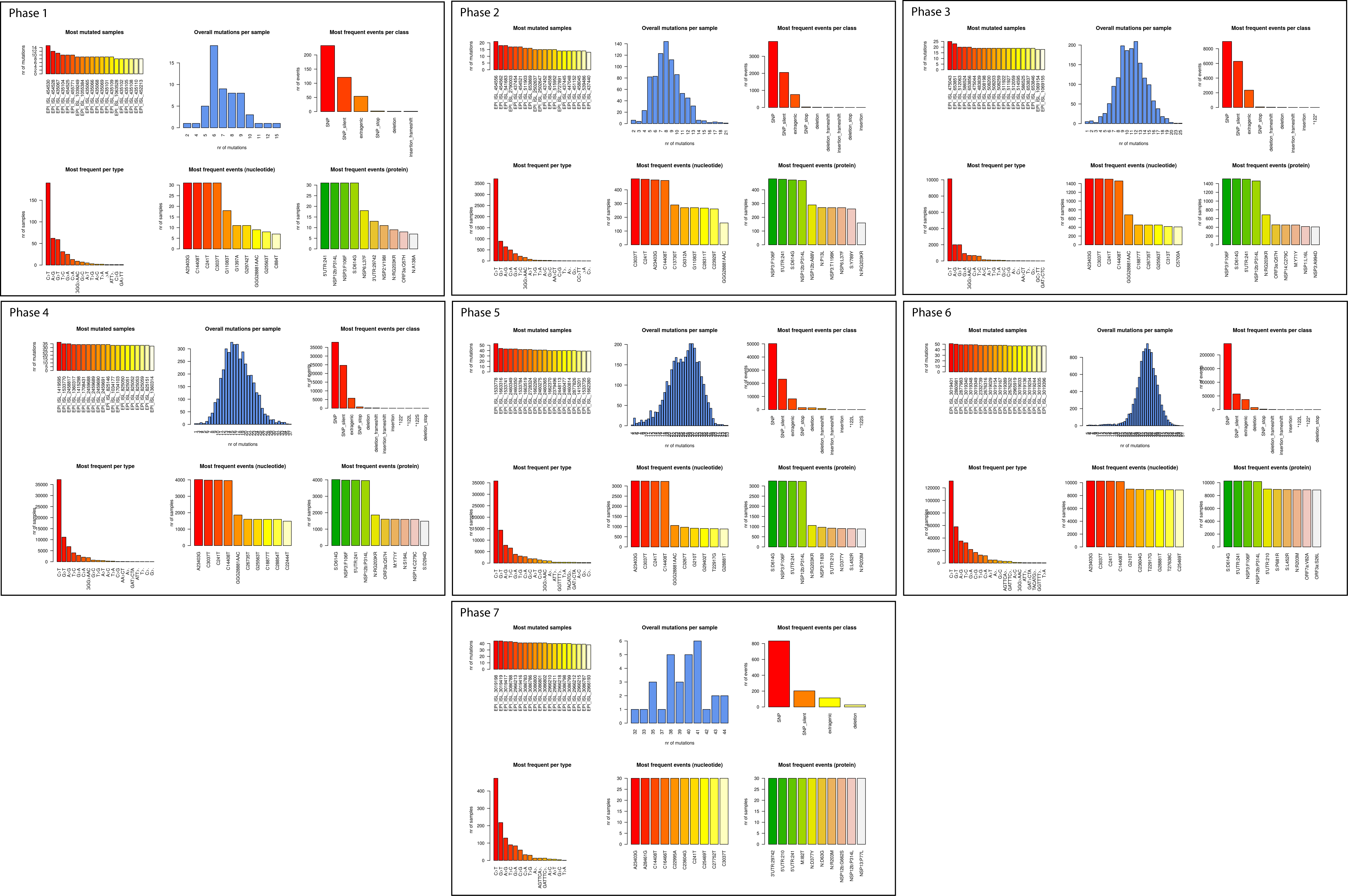
